# Accounting for the life history structure of fitness in tests for adaptive reproductive acceleration

**DOI:** 10.64898/2026.09.10.750743

**Authors:** Stacy Rosenbaum, Anup Malani

## Abstract

A prominent developmental plasticity theory proposes that early life adversity accelerates life history trajectories so organisms can maximize fitness in challenging environments. Standard tests of this hypothesis rely on empirical predictions about observable life history variables (e.g., age at first birth), because the proposed internal processes cannot be directly measured. However, the observable proxies are themselves mechanically linked, and regressions that ignore this may conflate causal effects with mechanical relationships among proxies. Here, we develop theory that explicitly justifies some existing empirical tests but rejects others. We integrate an accounting model that incorporates age at first birth, interbirth intervals, and lifespan with an optimization model, to derive the mathematical form that tests of the hypothesis should take. We apply these tests to data from wild female baboons, using early life rainfall as an exogenous proxy for early adversity so that estimates can plausibly be interpreted causally. The accounting model explains >98% of the variation in lifetime reproductive success among females that reproduced. However, our theory-derived tests find no evidence for the reproductive acceleration hypothesis. Our study illustrates the value of formal theory for causal inference in developmental plasticity research, where intertwined measurable variables may be obscured by strictly verbal hypotheses.

## 1 Introduction

The nature, degree, and consequences of phenotypic plasticity (i.e., the ability of a single genotype to produce different phenotypes in response to different stimuli) are central puzzles in evolutionary biology (Pigliucci 2001; West-Eberhard 2003). Developmental plasticity is a particularly intriguing type of plastic response because in some cases, the developmental input and the phenotypic response to it are meaningfully separated in time (Beldade et al. 2011). Whether such phenotypic adjustments are adaptations (i.e., under positive selection) (Ghalambor et al. 2007; Auld et al. 2009), and if so what mechanisms produce them (Lafuente and Beldade 2019; Uller 2008), have long been active areas of investigation.

One potential phenotypic adjustment organisms might make is altering the timing of life history events in response to developmental cues (Draper and Harpending 1982; Belsky et al. 1991; Bateson et al. 2014). The original (and still most widely invoked) logic is that organisms that face early life adversity should switch from investing in growth to investing in reproduction earlier, and/or reproduce at a faster rate than they otherwise would have, to maximize lifetime fitness (Chisholm et al. 1993). Since poor developmental environments could compromise adult somatic quality and survival, reproducing earlier or faster would “make the best of a bad job” (Belsky et al. 2015; Nettle and Bateson 2015). Thus, there should be selection for accelerated life histories under these conditions. This idea goes by various names, including internal predictive adaptive response and somatic-state based plasticity in the behavioral and evolutionary ecology literature (Bateson et al. 2014; Nettle and Bateson 2015), and the psychosocial acceleration, stress acceleration, and reproductive acceleration hypothesis in psychology (Ellis 2004; Belsky 2012). Here, we will refer to this phenomenon as reproductive acceleration.

### 1.1 Existing tests for accelerated reproduction

The reproductive acceleration hypothesis generates multiple testable predictions about the relationships among early life adversity, lifespans, reproductive pacing, and reproductive success. Variations of these predictions appear in myriad forms throughout the literature (e.g., Chisholm et al. 1993; Monaghan 2008; Ellis et al. 2009; Hayward et al. 2012). They are comprehensively assembled in Nettle and Bateson (2015), which outlines several complementary empirical predictions designed to evaluate whether adversity accelerates reproduction; whether this acceleration is specifically adaptive for individuals who experience adversity; and whether the benefits are greatest for individuals with shorter lifespans, since shortened lifespan is a key pathway through which adversity is thought to act. These predictions are as follows:

#### Prediction: Early life adversity causes shorter lifespans

To determine if this prediction is supported, **Test 1** regresses lifespan against some quality or state of the early life environment (we will assume that lower values mean more adversity). A positive association between the two is consistent with two things: that adversity damages somatic state and is likely to lead to an earlier death, and that adversity-exposed organisms have a shorter window in which to maximize lifetime reproductive success. The relationship can be read two ways. Early adversity might lower lifespan directly, or indirectly through accelerated reproduction that carries somatic costs.

#### Prediction: Early adversity causes accelerated reproduction

To determine if this prediction is supported, **Test 2** regresses some putatively adaptive phenotype (e.g., earlier age at first birth, shorter interbirth intervals, or some combination of the two) against early life environment. If the hypothesis is true, individuals who experienced early adversity should have accelerated reproduction relative to those who did not.

#### Prediction: The fitness benefits of accelerated reproduction depend on early adversity

To determine whether this prediction is supported, **Test 3** regresses lifetime reproductive success (*LRS*) against the product of early life environment and phenotype. A positive interaction would be evidence of an adaptive response to adversity; the absence of an interaction would imply that accelerating reproduction is simply beneficial in general.^1^

#### Prediction: The fitness benefits of accelerated reproduction depend on lifespan

**Test 4** parallels Test 3 but substitutes lifespan for adversity (i.e., it regresses *LRS* against lifespan interacted with phenotype), predicated on the theory that shortened lifespans mediate the effect of adversity on fitness. A positive interaction would indicate that accelerating reproduction is particularly beneficial for individuals with shorter lifespans.

This framework is attractive because it offers a full set of empirical predictions that can be evaluated using observational data. A paper on wild female baboons provided what is likely the most comprehensive empirical test to date (Weibel et al. 2020), but individually, tests of these predictions have a much longer history. They have been tested either in part or in full across a wide range of taxa, with mixed results (reviewed in Bonapersona et al. 2019; Ding et al. 2024).

Here, we build on this framework by addressing two aspects that would benefit from additional formal treatment. First, while the question of interest is causal (i.e., is early adversity causing accelerated reproduction?), the actual application relies on associational regressions that cannot separate the proposed causal mechanism from alternative explanations. Second, the tests themselves were developed from verbally articulated intuitions rather than derived from a formal model, and may not correctly reflect the mathematical relationships among life history variables and fitness. We first discuss the causal inference problem, then develop the theory necessary to derive what these tests of the reproductive acceleration hypothesis should look like given the mathematical relationships that govern how fitness is connected to lifespan and reproductive pacing.

### 1.2 The causal inference problem in developmental plasticity research

While the theories that motivate developmental plasticity research are fundamentally causal, a core challenge is that causality is difficult to directly test. To do so, we would have to observe the same organism in two different states of the world: one where they experienced a high-quality early life environment, and one where they experienced the opposite. An approximation of this can sometimes be achieved, most often with non-mammals (e.g., “split clutch” field experiments with birds and reptiles: (Warner 2014)). While the most rigorous versions occur in tightly controlled lab settings (with, e.g., genetically identical rodents), such studies often have questionable external validity. In order to maintain ecological realism, we often rely instead on indirect tests to infer the nature and strength of the causal relationship(s) among developmental environments, phenotypes, and fitness outcomes.

Testing a causal claim requires solving two separate problems. The first is observability: to evaluate the effect of variable *x* on outcome *y*, we need to measure both. Many hypotheses in developmental plasticity research — including reproductive acceleration — concern environmental effects on internal somatic processes (e.g., physiology, gene expression, energy allocation). These internal processes (*y*) are hard to measure, so we must rely on an observable proxy *z*. In order to ensure that our proxy is capturing what we intend, we need a model that derives *z* from *y* that is explicit about its assumptions.

For reproductive acceleration, the purported proxy *z* is the timing of life history events. If early adversity (*x*) causes some physiological adjustment (*y*) that results in a shift in energy allocation away from growth and maintenance and toward reproduction, this should produce a predictable pattern of lower age at first birth and shorter interbirth intervals. An important complication is that the observable variables involved in these tests (age at first birth, interbirth intervals, lifespan, and lifetime reproductive success) are themselves mechanically linked. Without a model that specifies those links explicitly, regressions risk conflating the causal effect of adversity with mechanical relationships among life history variables.

The second problem that needs to be solved is exogeneity. The variation in *x* must not be confounded by some quality of the organism itself. For this reason, ecological variables such as rainfall and temperature are useful, because organisms themselves do not influence how much it rains or how cold it is. Social variables (e.g., dominance rank, social isolation) are often of interest as well because of their strong correlations with health, reproduction, and longevity (Snyder-Mackler et al. 2020), but it is challenging to separate out the effects of social environments themselves because they are likely to be at least in part determined by an organism’s own qualities (e.g., its genes or behavior) (Malani et al. 2023). An unambiguous causal estimate requires environmental variation that is not influenced by the organism itself.

Here, we use life history accounting and an evolutionary optimization model to derive what the predicted pattern of reproductive pacing should look like if the reproductive acceleration hypothesis is supported. We use this to motivate empirical tests that are directly evaluating the hypothesis rather than the mechanical relationships among life history variables. We then use our theory-derived tests to determine whether an exogenous proxy for early life adversity (rainfall, or lack thereof) causes reproductive acceleration in wild female savannah baboons monitored since 1971 by the Amboseli Baboon Research Project (Alberts and Altmann 2012). This two-problem framing — observability first, then identification — gives a general template for causal inference in developmental plasticity research. It does not remove the assumptions that such inference requires, but it does make them explicit.

## 2 An accounting model of lifetime reproductive success

The internal developmental response we ultimately care about that could causally connect early adversity to accelerated reproduction cannot be directly observed. We instead need to infer it from observable variables: age at first birth (*A*), interbirth intervals (*IBI*), lifespan (*L*), and lifetime reproductive success (*LRS*). It is useful to start by summarizing the broader causal structure we envision (Figure 1). Some exogenous environmental variable *e* affects an organism’s lifespan directly (e.g., through early nutritional deficits that compromise adult somatic quality) and possibly indirectly through its effects on the pace of reproduction (*A* and *IBI*), which themselves shape lifespan via reproduction–maintenance trade-offs. In our figure, the unobserved variable *U* represents common causes of *A, IBI*, and *L* that we do not directly measure — for example, prenatal effects, maternal effects, and other unobserved environmental factors. The internal developmental response is not shown; it lies on the pathway from *e* to *A* and *IBI*.

**Figure 1.**
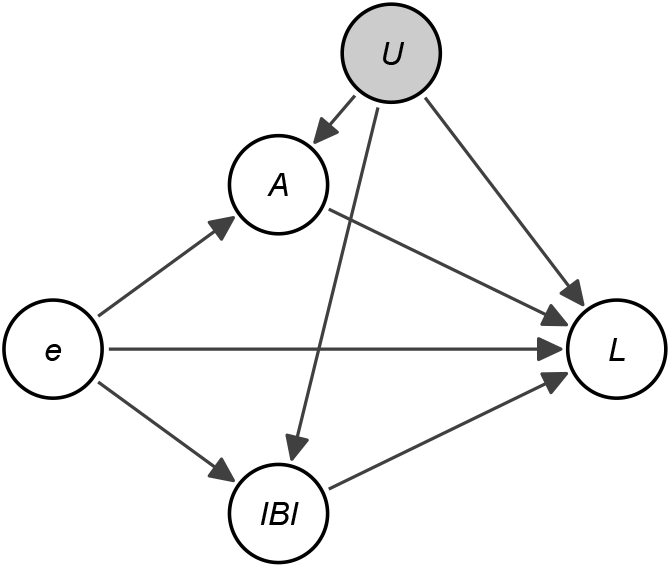
Hypothesized causal structure for our model: exogenous environmental variable *e* affects observed age at first birth (*A*), length of interbirth intervals (*IBI*), and lifespan (*L*) both directly and indirectly via reproduction– maintenance trade-offs. *U* (gray, unobserved) represents common causes of *A, IBI*, and *L* that we do not directly measure. Lifetime reproductive success is not shown here; it is mechanically determined by *A, IBI*, and *L* via the accounting model developed in Section 2 (see also Figure 2).

However, before we can address the causal question of interest, we first need to understand how the observable variables relate to one another. In this section, we develop an “accounting model” that makes that relationship explicit. It connects *A, IBI*, and *L* to *LRS* deterministically, on top of this causal structure.

Because we will ultimately execute our test of the reproductive acceleration hypothesis using data from baboons, we will rely on baboons for the examples, data, and discussion going forward. Consider a female baboon who has her first offspring at age *A*, goes on to have *n* more offspring, and eventually dies at age *L*. Her *LRS* is the total number of offspring: *LRS* = *n* + 1. We will denote each of her individual interbirth intervals as *I*_*i*_ and her average interbirth interval as *IBI*:

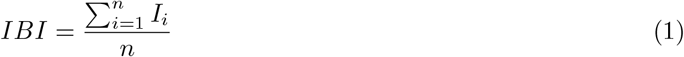

Our hypothetical baboon’s lifespan can be broken down into three periods: the time before her first birth (i.e., *A*), the time she spent reproducing (which is the sum of her interbirth intervals, or ∑_*n*_*I*_*i*_), and the period *E* between the birth of her final offspring and her death:

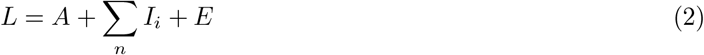

In most cases, *E* is probably shorter than a female’s average *IBI*. Since few animal species routinely have meaningful post-reproductive lifespans (Ellis et al. 2018), the female likely would have given birth again had she survived for another full interbirth interval. For simplicity, we will assume that the probability of dying is constant over time, so that *E* is approximately half of the female’s average *IBI*: *E* ≈ 0.5 × *IBI*.

We now need to rewrite the lifespan equation in terms of *LRS* and *IBI* rather than individual interbirth intervals by making two substitutions. First, since the sum of the individual interbirth intervals equals the number of intervals times the average interval, we can replace ∑_*n*_*I*_*i*_ with *n IBI*. Second, because the female has *n* + 1 total offspring (her first, plus *n* more), and *LRS* = *n* + 1, it follows that *n* = *LRS* − 1. Substituting these into the lifespan equation along with our approximation for *E* gives us the following:

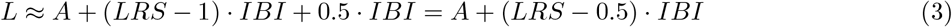

We can then rearrange to solve for *LRS*:

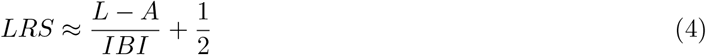

This is our basic reproductive accounting equation for *LRS*. Three quantities determine lifetime reproductive success: lifespan, age at first birth, and average interbirth interval. Note that these key variables are used in the previously described tests for accelerated reproduction in Section 1.1, as well as in our causal visualization in Figure 1. Our accounting model specifies exactly how they should relate to one another. Figure 2 illustrates the structural relationships encoded in Equation (4): *LRS* is mechanically determined by *A, IBI*, and *L*. These are accounting relationships, not causal claims, and are drawn with dashed arrows to distinguish them from the causal structure shown in Figure 1.

**Figure 2.**
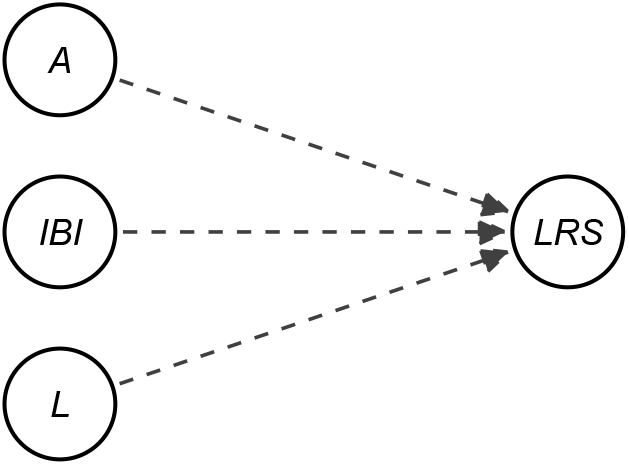
Accounting structure of lifetime reproductive success. As in Equation (4), *LRS* is mechanically determined by age at first birth (*A*), interbirth interval (*IBI*), and lifespan (*L*). The dashed arrows indicate algebraic/accounting relationships, not causal claims.

### 2.1 Empirical validation of the accounting model

It is important to empirically validate the assumption that our proposed accounting model explains *LRS* in the real world, where there are many factors that might affect the *LRS* of any given female. Therefore, we applied the equation to baboon life history data published by Weibel and colleagues (2020). These data, from *n* = 110 wild female baboons with complete reproductive histories, contain all of the necessary information to test the fit of our equation. Following Weibel and colleagues, *LRS* is defined as the total number of live offspring a female gave birth to^2^, and *A* is defined as her age at first live birth. *IBI* is the mean number of days between consecutive live births in which the first offspring of the two survived to 70 weeks (the average age of weaning for Amboseli baboons (Altmann 1998)). Further details about the data can be found in Section S1 of the supplementary materials and in Weibel et al. (2020).

By construction, this validation only includes females who had at least one *IBI* (i.e., gave birth to at least two live offspring). This is a necessary restriction since the accounting model includes *IBI*, which is undefined for females with only a single birth. We discuss this scope restriction, and how the model could and could not be extended to females with *LRS <* 2, in Section S2 of the supplementary materials.

This simple accounting model does a remarkably good job of explaining these baboon data. If we plot females’ actual *LRS* against our mathematical approximation for *LRS*, there is a very strong correlation between the two (Figure 3). We can also demonstrate this by rearranging Equation (4) so that lifespan (*L*) and age at first birth (*A*) appear on opposite sides, so that it can be estimated as a regression:

**Figure 3.**
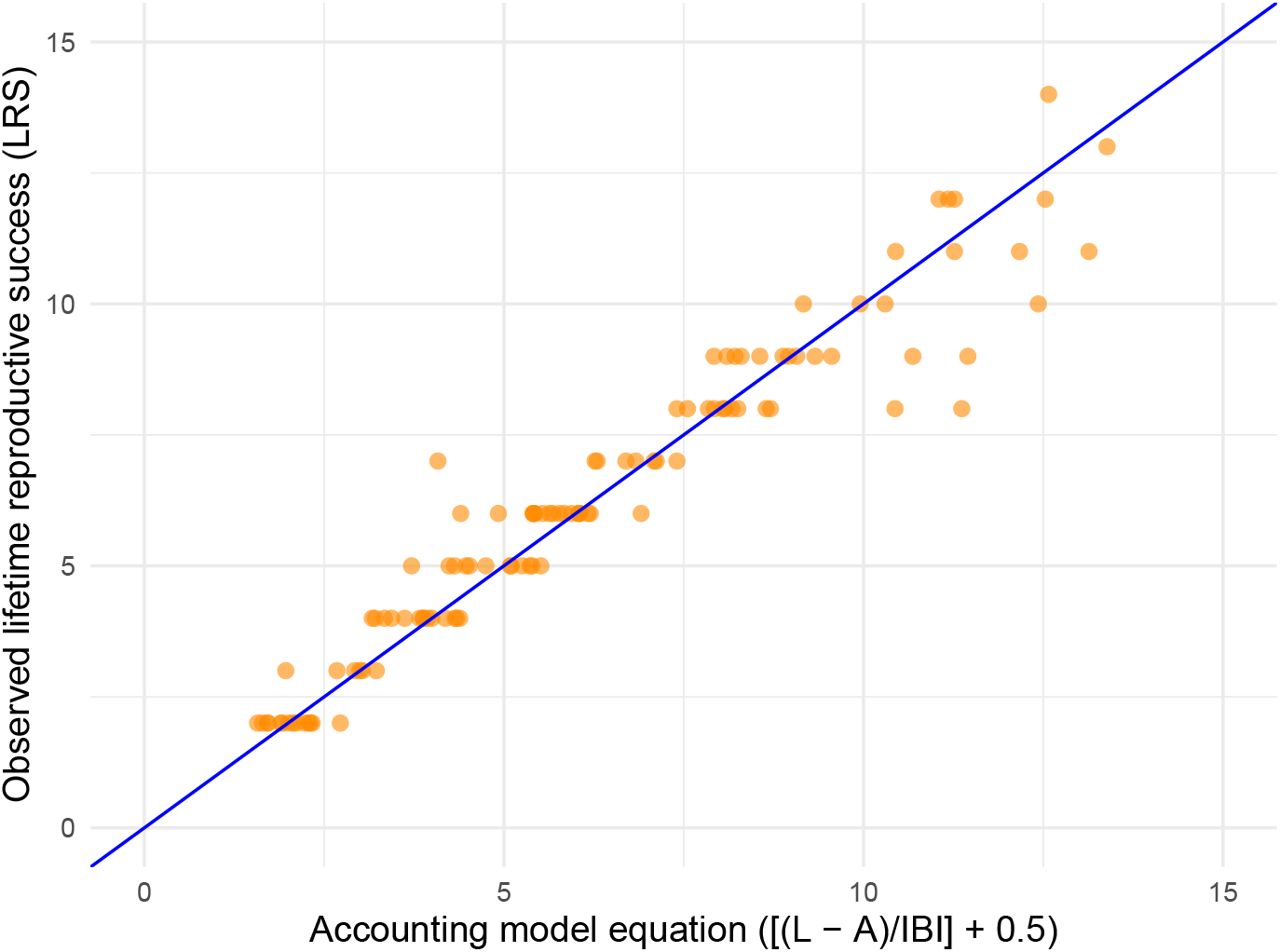
Predicted versus actual lifetime reproductive success in female savannah baboons monitored by the Amboseli Baboon Research Project. Each orange dot is a single female with a completed, observed reproductive history and at least one measured interbirth interval (*n* = 110). The x-axis shows the predicted lifetime reproductive success derived from the accounting model (Equation 4), and the y-axis plots observed female reproductive success. The solid blue line is the 1:1 fit line.

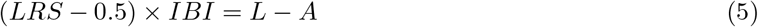

The accounting model’s fit to these baboon data is nearly exact. The regression coefficients are very close to what Equation (4) predicts, and the model’s *R*^2^ is very high regardless of whether the data are untransformed (*R*^2^ = 0.98) or log-transformed (*R*^2^ = 0.99; Table 1). Thus, the accounting model explains virtually all of the variance in *LRS*. This confirms that in this sample of female baboons, *LRS* is indeed mechanically determined by (*L, A, IBI*) in the manner that is specified in the accounting model. This is useful because we now know that a regression involving *LRS* and these life history variables must respect this basic mathematical structure.

**Table 1:**
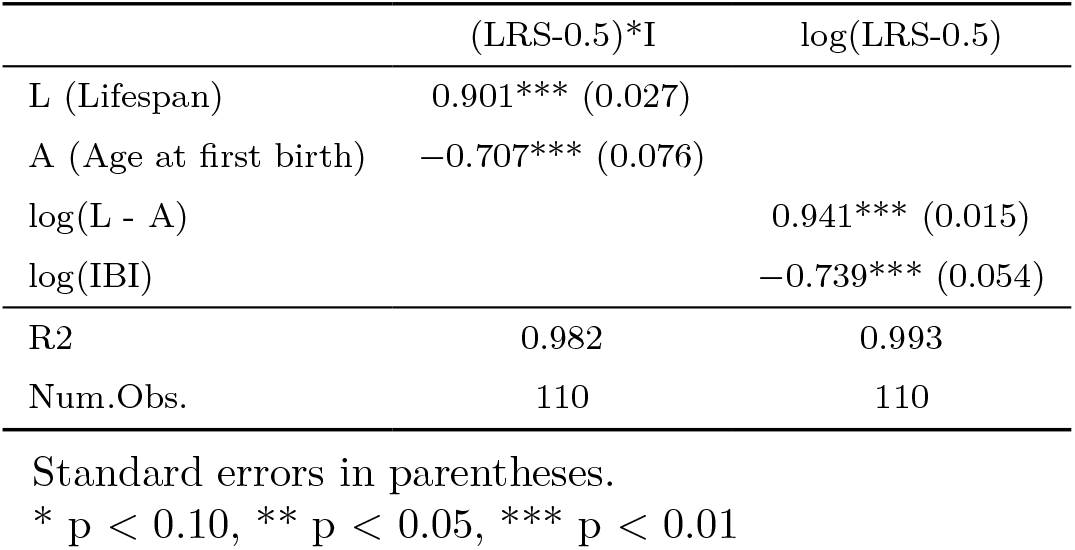
Accounting regression results.

## 3 A biological optimization model

Having shown that an accounting model of *LRS* fits the data well, we now need a theory that connects early adversity to changes in reproductive pacing. The accounting model tells us how *LRS* is mechanically generated, but not why females “choose” particular values of *A* and *IBI*. For that, we need an optimization model.

### 3.1 The trade-off problem

In any given environment, baboons should choose the optimal *A* and *IBI*, because we expect natural selection to produce phenotypes that are fitness maximizing (Parker and Smith 1990). Therefore, we first need to derive what that optimum looks like generically and then compare the optima in environments with and without early life adversity.

Earlier *A* and shorter *IBI* would, all else being equal, benefit any female regardless of her early environment (Nettle and Bateson 2015; Weibel et al. 2020). If reproducing earlier or faster was costless, then every female would push reproductive pacing to the limits imposed by basic physiological constraints (e.g., *A* below a certain age is impossible because the reproductive system is not yet functional). The fact that observed reproductive pacing usually sits above such physiological “floors” implies a cost. The obvious potential cost of an earlier *A* and/or shorter *IBI* (which both increase *LRS*) is a shorter lifespan (which decreases *LRS*), consistent with growth/reproduction/maintenance trade-offs that are fundamental to life history theory (Stearns 1992).

### 3.2 Optimal reproductive pacing, holding environment constant

We assume that organisms maximize *LRS* as approximated by the accounting model (Equation (4)), that they choose *A* and the length of *IBI*, and that lifespan depends on these choices.

Suppose that lifespan is an increasing and concave function of both *A* and *IBI*. That is, females who delay reproduction and space their births more widely live longer (∂*L/*∂*A >* 0, ∂*L/*∂*IBI >* 0), but the lifespan gains from further delay or wider spacing diminish as the values of each increase (∂^2^*L/*∂*A*^2^ *<* 0, ∂^2^*L/*∂*IBI*^2^ *<* 0). For simplicity, we also assume that the effect of *A* on lifespan does not depend on *IBI*, and vice versa (∂^2^*L/*∂*A*∂*IBI* = 0).

To illustrate the trade-off, we generated simulated data for 150 females using a concave (logarithmic) function relating lifespan to *A* and *IBI* length, with normally distributed individual variation (Figure 4). In our xssimulation, each female has only a single *IBI*. We also extended the possible range for both variables relative to what is actually observed in baboons, because in practice the observed range, especially of *A*, is narrow enough that the curvature would be difficult to see.

**Figure 4.**
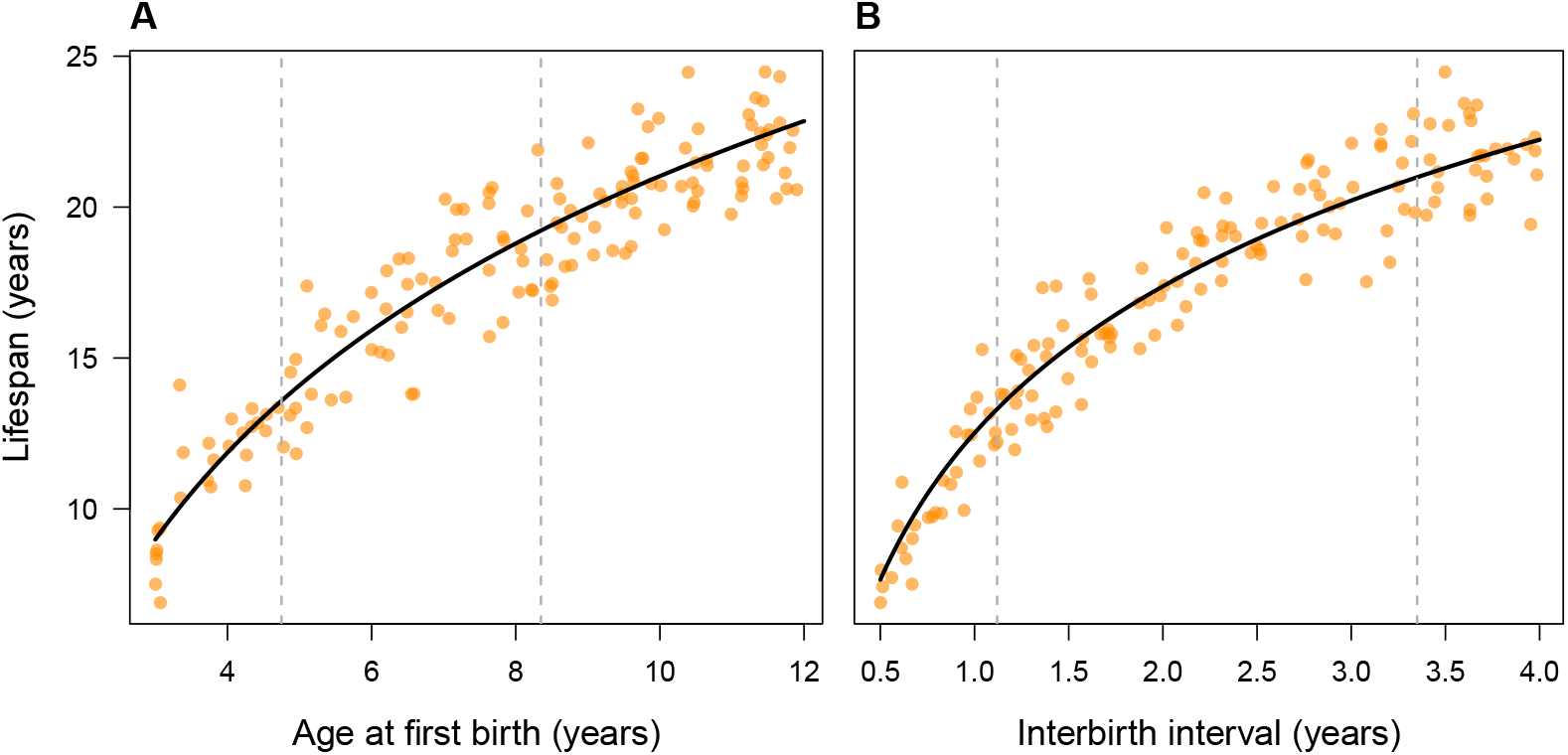
The assumed reproduction–lifespan trade-off. Each point represents a simulated female with a single interbirth interval. **Panel A:** Lifespan increases with age at first birth but with diminishing returns (concave). **Panel B:** Lifespan increases with longer interbirth intervals but with diminishing returns (concave). In both panels, the solid curve shows the underlying trade-off function, the orange points show individual simulated females with random variation, and vertical gray lines mark the observed range in the Amboseli baboon data. The x-axis ranges are extended beyond the observed range to make the concavity visible.

### 3.3 Conditions for optimal reproductive pacing

If females are indeed optimizing, we can derive the conditions that must hold at the *LRS*-maximizing values of *A* and *IBI*.

#### Age at first birth

A female should delay her first birth until each additional year of delay adds exactly one year to her lifespan:

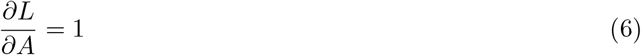

Beyond that point, the lifespan gain from further delay is less than the reproductive year lost while waiting.

#### Interbirth interval

A female should space her births until the lifespan gained from longer intervals equals the reproductive output lost across all her remaining intervals:

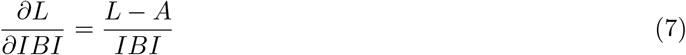

The left side of this equation is the lifespan gain from a longer *IBI*; the right side is her current reproductive rate (total reproductive years divided by *IBI*), which is the *LRS* units she would forego by slowing down. The concavity conditions on lifespan described above ensure that these conditions are sufficient for an optimum.

### 3.4 How early life adversity affects optimal choices

Our primary goal is to understand why early life adversity would cause an organism to change their choice of *A* and/or *IBI* length. Specifically, we ask: given that animals are already optimizing their choice of *A* and *IBI* (and thus their *LRS*) via evolutionary processes, what happens to the optimal choice when an animal experiences early adversity? We assume that lifespan depends not only on reproductive pacing but also on early life adversity, *e* (where lower values of *e* mean more adversity). In other words, we propose that *L*(*A, IBI, e*), as depicted in Figure 1. We make three assumptions:

1. Females who do not experience early life adversity live longer, all else being equal (∂*L/*∂*e >* 0).
2. The lifespan benefit of delaying first birth is smaller for females who experienced early adversity (∂^2^*L/*∂*A* ∂*e >* 0). In other words, a female who faced early adversity gains less additional lifespan from waiting an extra year to reproduce than a female who did not.
3. Similarly, the lifespan benefit of lengthening interbirth intervals is smaller for females who faced early adversity (∂^2^*L/*∂*IBI* ∂*e >* 0).

Assumptions 2 and 3 are key, as they say that early adversity flattens the trade-off curves depicted in Figure 4, reducing the lifespan-lengthening payoff of delayed or slower reproduction.

Given the optimality conditions (Equations (6) and (7)), this implies that females who experienced adversity (lower *e*) should optimally choose a younger age at first birth (*A*^∗^) and shorter interbirth intervals (*IBI*^∗^):

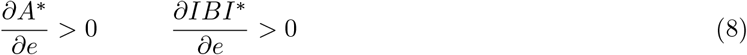

In other words, adversity reduces the lifespan payoff of delaying reproduction, so the optimum that balances costs and benefits shifts to lower values of *A* and shorter *IBI*.

### 3.5 Deriving a regression from the optimization model

To test the above model empirically, we need to estimate the relationship between lifespan and its proposed determinants (*A, IBI*, and *e*) using a regression. The challenge is that we do not know the exact mathematical form of the lifespan trade-off function. We have hypothesized, based on tenets of life history theory, that it is increasing, concave, and flattened by adversity. The choice of empirical specification shapes regression results alongside the theory being tested, so we want a form that is flexible enough to let the theory do the work. We therefore approximate *L*(*A, IBI, e*) using a second-order Taylor expansion:

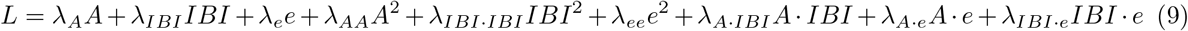

A Taylor expansion approximates an unknown smooth function as a polynomial, in this case as a sum of linear, squared, and interaction terms of *A, IBI*, and *e*. The second-order version captures overall slopes, curvature, and pairwise interactions among the inputs, which together fit a wide range of underlying functional forms (e.g., logarithmic, exponential, power-law) without committing to a particular one in advance. The Taylor expansion is therefore flexible enough to capture the properties we have hypothesized for the trade-off if they are present in the data, without specifying their exact form. We use the second-order version because it is the simplest one that can capture the hypothesized diminishing returns and adversity-by-pacing interactions; going higher (cubic terms, three-way interactions) would estimate many more parameters than our sample size can support.

Equation (9) can be estimated as a standard regression. Each *λ* coefficient captures a different aspect of how lifespan responds to reproductive pacing and early adversity. The terms fall into three groups:

- **Direct effects** (*λ*_*A*_, *λ*_*IBI*_, *λ*_*e*_): the baseline relationships — how lifespan changes with *A, IBI*, and *e*, respectively.
- **Curvature terms** (*λ*_*AA*_, *λ*_*IBI*·*IBI*_, *λ*_*ee*_): these capture the diminishing returns illustrated in Figure 4. Negative values indicate that the lifespan gains from delaying age at first reproduction, or lengthening *IBI*, taper off.
- **Interaction terms** (*λ*_*A·IBI*_, *λ*_*A·e*_, *λ*_*IBI·e*_): these terms capture how the effect of one variable depends on another.

By the accounting model (Equation (4)), *L* − *A* ≈ (*LRS* − 0.5) ·*IBI*, so the same regression can also equivalently be estimated with (*LRS* − 0.5) ·*IBI* as the outcome. This is asking whether adversity flattens the pacing–fitness trade-off rather than the pacing–lifespan trade-off.

These equations are the regressions we need to run to test whether early adversity leads to faster reproductive pacing. The key coefficients are *λ*_*A·e*_ and *λ*_*IBI·e*_, which correspond directly to assumptions 2 and 3 from Section 3.4. If *λ*_*A·e*_ *>* 0, the lifespan benefit of delaying first birth is smaller for females who experienced adversity; if *λ*_*IBI·e*_ *>* 0, the same is true for lengthening *IBI*. Positive values for these coefficients would indicate that early adversity flattens the trade-off curves, which is how the proposed adaptation operates.

## 4 Mapping the theory to prior tests

In this section, we use the accounting and optimization models to revisit each of the four tests outlined in Section 1.1. For each, we ask: what should the regression look like once we account for the structure of the underlying life history variable relationships, and does the test remain meaningful? The take-home message is that our framework changes the relationships among the tests, in some cases substantially. Rather than four independent regressions, the tests reduce to two distinct empirical exercises, with one additional test that can be eliminated by the accounting model itself.

### 4.1 Relationships among the tests

#### Test 1: Relationship between lifespan and early life adversity

Original Test 1 has two possible interpretations, which are indistinguishable without the explicit functional form for lifespan developed in Equation (9). The first interpretation is that adversity lowers lifespan directly (∂*L/*∂*e >* 0). The second is that adversity lowers lifespan indirectly, via somatic costs imposed by accelerated reproduction. The Taylor expansion nests both possibilities. The direct effect interpretation corresponds to *λ*_*e*_ and *λ*_*ee*_, and the indirect effect interpretation to *λ*_*A·e*_ *>* 0 and *λ*_*IBI·e*_ *>* 0, the interactions that capture trade-off flattening. The regression implementing Test 1 is therefore not simply *L* ~ *e*; it is the full Taylor expansion with *L* as the outcome.

#### Test 2: Relationship between accelerated reproduction and early life adversity

The optimization model predicts that reproductive pacing should shift with adversity (Equation (8)). Original Test 2 tests this directly. Females who faced more early adversity should optimally choose a younger age at first birth and shorter *IBI*s, so ∂*A*^∗^*/*∂*e >* 0 and ∂*IBI*^∗^*/*∂*e >* 0. This is tested by regressing *A* on *e* and *IBI* on *e* separately. This regression structure is unchanged from the originally proposed test. Our framework contributes the theoretical grounding for the predicted signs.

#### Test 3: Relationship between lifetime reproductive success, early adversity, and accelerated reproduction

Original Test 3 is simply a re-parameterization of Test 1 once the accounting model is imposed on the left-hand side. Tests 1 and 3 rely on the same underlying variation in the data; the only difference is whether the accounting structure is imposed before the regression is estimated.

#### Test 4: Relationship between lifetime reproductive success, lifespan, and accelerated reproduction

Original Test 4 should be zero, given the accounting model. It asks whether accelerated reproduction is specifically beneficial for females with shorter lifespans, operationalized as a positive *L*×*p* interaction in an *LRS* regression. But the accounting model leaves no room for an *L*×*p* interaction. The accounting equation *LRS* ≈ (*L* − *A*)*/IBI* + 0.5 combines lifespan with age at first birth only as a difference, and with interbirth interval only as a ratio. Neither operation creates an *L*×*p* interaction, for either measure of *p*. This equation captures >98% of the variation in *LRS* in the data set we tested (Table 1). Any *L*×*p* effect a regression detects would therefore reflect residual variance, including variance caused by measurement error, rather than the biological mechanism of interest. A significant result would not constitute evidence for reproductive acceleration.

Taken together, these points demonstrate what is gained by deriving tests from an explicit model. In this use case, it allows us to separate out the direct and indirect pathways by which early adversity could shorten lifespan; helps us avoid the pitfalls of conducting two separate tests on the same underlying variation; and most importantly, clarifies which relationships among life history variables actually speak to the hypothesis. None of these follow from the four verbally derived tests on their own.

## 5 Testing the reproductive acceleration hypothesis in baboons

Here, we apply the new tests to data on wild female savannah baboons monitored by the Amboseli Baboon Research Project (ABRP), which has collected demographic and life history information on individually identified baboons on a near-daily basis since 1971 (Alberts and Altmann 2012). Most of these data were previously published (Weibel et al. 2020). They are useful because they contain information about all of the life history variables necessary to run our tests, as well as information about different sources of early life adversity that the ABRP has previously identified for baboons (Tung et al. 2016). More information about the baboons, the adversities they face, and the data that are specifically used for these analyses can be found in Sections S1 and S3 of the supplementary materials, as well as in Tung et al. (2016) and Weibel et al. (2020).

We focus on one particular source of early adversity: the amount of rain experienced in the first year of life (as total mm/year). Lower rainfall is a proxy for greater adversity, because the baboons live in a highly seasonal environment where food availability is correlated with annual rainfall (Aduma et al. 2018; Rosenbaum et al. 2025). Most prior papers on this population have operationalized early life rainfall as a binary variable; i.e., the baboon in question either did or did not experience drought when they were young. However, the ABRP generously provided us with the continuous rainfall data that underlies the binary drought indicator in Weibel and colleagues’ (2020) analysis, so we use this continuous measure here. Rainfall is measured daily at the ABRP field site using a single rain gauge and summed across each individual female’s first year of life (Weibel et al. 2020). The rest of our data are identical to what was used in the original publication.

We choose to focus on rainfall specifically because it is a source of adversity that is exogenous to the baboons, which makes it a useful source of variation for our causal-inference strategy^3^. No feature of the baboons themselves influences how much it rains, whereas other sources (e.g., maternal loss, being born to a low-ranking mother) are more likely connected to baboon genetics or behavior in ways that complicate interpretation (Malani et al. 2023). Since rainfall is randomly distributed across baboons, it approximates a natural experiment.

It is important to clarify what this strategy does and does not deliver. Rainfall in the first year of life plausibly affects lifespan and reproductive pacing through several biological pathways (e.g., direct nutritional effects, density-mediated competition, maternal condition). The reproductive acceleration hypothesis is agnostic about which of these produces the adversity effect. What matters for the hypothesis is whether adversity, however it acts, reshapes the reproduction–lifespan trade-off. The Taylor expansion addresses this question: it separates whether rainfall affects lifespan directly (*λ*_*e*_, *λ*_*ee*_) from whether it flattens the pacing–lifespan trade-off (*λ*_*A·e*_, *λ*_*IBI·e*_), with the latter interactions being the structural “signature” that the optimization model predicts. This is a meaningful advance over a simple regression of pacing on adversity, which cannot distinguish the two. However, it does not give us mechanism-specific information about what biological pathways adversity is operating through.

The theory in Equation (8) predicts positive coefficients on rainfall. Better environmental conditions in early life should raise optimal *A*^∗^ and lengthen optimal *IBI*^∗^, and should produce positive interaction coefficients (*λ*_*A·e*_ and *λ*_*IBI·e*_) in the Taylor expansion. Sample sizes for our analyses vary because not all data are available for all individual baboons, and are reported alongside the relevant results. We excluded control variables used in some models in Weibel and colleagues’ original paper after verifying that their inclusion did not change our results (Supplementary materials Section S4). Note that in the Taylor expansion, *A* and *IBI* are inputs for lifespan (Figure 1), not just variables we are including as controls to isolate a direct effect of *e* on *L*. For transparency and to evaluate the overall relationship between rainfall and lifespan without modeling the trade-off function at all, we also report the results of a simple regression *L* ~ *e* alongside the Taylor expansion results. This version does not rely on the assumption that there are no unobserved factors that jointly influence reproductive pacing and lifespan.

### 5.1 Results

#### 5.1.1 The reproduction–lifespan trade-off

We find no evidence that rainfall in the first year of life flattens the reproduction–lifespan trade-off in the Amboseli baboons. We estimate Equation (9) with lifespan as the outcome and test the key coefficients *λ*_*A*·*e*_ and *λ*_*IBI*·*e*_. The model explains a modest amount of lifespan variation (*R*^2^ = 0.12, *n* = 99), but the coefficients on the adversity-by-pacing interactions are not statistically significant (*p* > 0.30 for both *λ*_*A*·*e*_ and *λ*_*IBI*·*e*_).

We find a similar null result when we reframe the question in terms of *LRS* rather than lifespan. Under the accounting equation, asking whether adversity flattens the pacing–lifespan trade-off is algebraically equivalent to asking whether it flattens the pacing–fitness trade-off, so we can swap (*LRS* − 0.5) *IBI* for lifespan as the regression outcome without changing the underlying biological question. The interaction terms *λ*_*A·e*_ and *λ*_*IBI·e*_ are non-significant (*p* > 0.43). Full regression results for the lifespan and *LRS* models are reported in Table S7 in the supplementary materials.

A simple *L* ~ *e* regression that does not include any life history pacing variables also shows no significant effect of rainfall on lifespan (*β* = 0.006, *p* = 0.15, *n* = 99), and has the same positive sign observed in the Taylor expansion (Table S7). This is important because a sign reversal between the two models would indicate that the rainfall coefficient obtained from the Taylor expansion was being affected by unobserved factors that influence both reproductive pacing and lifespan; we find no evidence that this is the case.

#### 5.1.2 Reproductive pacing and adversity

Our test of the relationship between reproductive pacing and early life adversity does not find support for the reproductive acceleration hypothesis (Table 2). Females who experienced more rain during their first year of life reproduced earlier than females who experienced less rain (*p* = 0.002), which is the opposite of the direction predicted by Equation (8). A one standard deviation increase in rainfall (~122 mm) corresponds to a ~45-day decrease in age at first birth. Given that the mean *IBI* in the baboon data is 678 days, this translates to an expected ~0.07 additional offspring over the course of a reproductive lifespan. The relationship between rainfall and *IBI* runs in the same wrong direction: females who experienced more rain had ~14-day *shorter* (rather than longer) *IBI*s per standard-deviation increase in rainfall, although this relationship was not statistically significant (*p* = 0.20).

**Table 2:** Reproductive pacing and rainfall regression results.

| Predictor | Age at first birth | Interbirth interval |
| --- | --- | --- |
| (Intercept) | 6.510*** (0.110) | 1.944*** (0.090) |
| Rainfall (per SD, 122 mm) | -0.124*** (0.039) | -0.039 (0.030) |
| R2 | 0.037 | 0.017 |
| Num.Obs. | 267 | 99 |

Standard errors in parentheses.
\* $p < 0.10$ , \*\* $p < 0.05$ , \*\*\* $p < 0.01$

## 6 Discussion

Testing reproductive acceleration — or any developmental plasticity hypothesis of similar form — requires solving two distinct problems. The first problem, observability, arises whenever the outcome is an internal phenotypic adjustment that cannot be directly assayed. When this occurs, tests must use theory-justified, observable proxies. Our accounting and optimization models together show precisely how reproductive pacing variables (*A, IBI*) can act as that proxy, with directional signs and mathematical forms specified by the theory. The reproductive acceleration hypothesis as a case study highlights the hazards of relying on verbal models instead of explicit theoretical justification. We risk testing predictions that do not actually align with our hypotheses, clouding our ability to arrive at the right answer.

The second problem is identification. Even with a good, theoretically grounded proxy, credible causal claims require exogenous variation in the variable of interest. Here, we used rainfall during the first year of life to provide that variation, since it is the least-confounded available proxy for early life environmental conditions experienced by wild baboons. Purely predictive associations between life history variables and non-exogenous sources of adversity (e.g., dominance rank, maternal condition) risk conflating the effect of adversity itself with the effect of “selecting in” to adversity because of the confounding effects of genes and behavior. Confounding can never be entirely eliminated when studying wild animals. However, we can make meaningful progress toward reliable, if cautious, causal inference by explicitly laying out the proposed relationships between all three links in the chain: the causal variable(s) of interest, the proxies we can actually measure, and outcomes like lifespan and fitness that are critical for understanding evolutionary processes. The causal structure in Figure 1 and the structural model that follows from it state the assumptions on which our estimates rest. While some are testable (Section 2.1), a key one (namely, that no unobserved factors jointly affect reproductive pacing and lifespan) is not, at least with the available data. Making assumptions explicit does not make them correct. However, when they are visible, readers can weigh them appropriately and future research can work toward testing them.

The Amboseli baboon data provide no evidence for the reproductive acceleration hypothesis, corroborating findings from an earlier paper (Weibel et al. 2020). Under our framework (i.e., the accounting model paired with the optimization model), reproductive acceleration would generate positive interaction coefficients indicating that early life adversity flattens the reproduction–lifespan trade-off. We find no evidence of this. In the limited instances where the point estimates from our models were meaningfully different than zero, they ran counter to the predictions of the reproductive acceleration hypothesis. Adversity *steepened* the trade-off curve, instead of flattening it.

Our findings join a growing body of work questioning the association between early adversity and reproductive acceleration for both theoretical and empirical reasons (e.g., Moorad et al. 2019; André and Rousset 2020; Del Giudice 2020; Frankenhuis and Gopnik 2023). On theoretical grounds, the null result we report is consistent with what life history theory predicts for a species like baboons. For accelerated reproduction to be adaptive, the fitness costs of an earlier death must be outweighed by the benefits of reproducing earlier or faster (Stearns 1992). This is most plausible in species where adults typically only reproduce once, and maternal care does not extend beyond lactation. Neither of these is true for baboons, or other long-lived, iteroparous mammals with extended maternal care. In species like these, a female who dies while she is still caring for dependent young leaves behind offspring who are themselves unlikely to survive (Zipple et al. 2021). Such offspring make up a larger proportion of total lifetime reproductive success for short-lived than for long-lived females. The relative fitness cost of dying with dependent young is therefore higher, not lower, for females who have short lives. Recent formal modeling demonstrates that when maternal and offspring fitness are closely coupled, there is selection for longer lives and slower reproduction, not the reverse (Zipple, Reeve, et al. 2024). Our null result is therefore consistent with theoretically grounded expectations.

While we did not find results in the predicted direction, we did find a plausibly causal effect of early life environment on reproductive pacing (again, consistent with Weibel and colleagues’ (2020) earlier analysis). Females who experienced more rain in their first year of life began reproducing earlier; there was a ~45-day reduction in age at first birth per standard-deviation increase in rainfall. The most parsimonious explanation is that early life resource abundance accelerates physical development, so earlier first births simply reflect faster maturation rather than more complicated, strategic adjustments of reproductive timing (Kuzawa et al. 2025). Further bolstering this interpretation, this effect is unlikely to have significant fitness implications given that on average it should amount to only an additional ~7% of one offspring.

Our analyses have several limitations. First, we make no claim that the Taylor expansion is the true mathematical form of the reproduction–lifespan trade-off. It is useful because it provides a flexible approximation that is consistent with the optimization model’s structural assumptions. But while it is locally accurate near the center of the data, it may miss curvature that is visible only at more extreme values of *A* or *IBI*. Two factors limit our power to detect trade-off flattening if it exists: the narrow observed range of *A* in the available baboon data, and the relatively modest sample size available for our Taylor expansion (*n* = 99). Second, while it is true that first-year rainfall is exogenous to baboon behavior, it plausibly affects lifespan, body condition, reproductive pacing, and fitness through multiple pathways that our analyses cannot distinguish. Finally, it would be especially useful to run these tests on species with meaningfully different life history characteristics (e.g., species in which most adults reproduce only once or in which maternal care is limited or absent), since these are conditions under which the reproductive acceleration hypothesis could in principle hold.

Sharpening causal claims in developmental plasticity research — and in ecology and evolution more broadly — requires both more rigorous theoretical specification and a wider empirical toolkit. Even when we cannot execute best-case tests of causality, there are mathematical steps we can take to improve the legitimacy of causal claims. In our analyses, for example, it is a step in the right direction that we can separate out direct effects of early adversity on lifespan, versus effects that operate through trade-offs with reproductive pacing. However, no single approach is enough on its own. Taking advantage of natural experiments (e.g., Petrullo et al. 2024; Siracusa et al. 2025), giving more weight to empirical findings that capitalize on exogenous variation, doing cross-population comparisons, and whenever possible performing experimental manipulations on wild or quasi-wild populations (e.g., Dantzer et al. 2022; Zipple, Vogt, et al. 2024) are all approaches that can help us retain strong external validity while making strides toward solving causal puzzles.

## 7 Acknowledgements

The authors would like to thank Zach Laubach, Jonas Wahl, Laura Dee, and Wei Perng for inviting us to participate in this special issue, as well as three anonymous reviewers for their helpful feedback. We are very grateful to the members of the Amboseli Baboon Research Project (ABRP), especially Chelsea Weibel, Beth Archie, Susan Alberts, and Jenny Tung, for sharing their data and for their advice and comments on early versions of the paper. The data used in this analysis were generated by the ABRP; funding information for the ABRP is available at https://amboselibaboons.nd.edu/acknowledgements/. This research was supported by the University of Michigan and the Barbara J. and B. Mark Fried Fund at the University of Chicago Law School.

## 8 Data and code availability

All data and code needed to replicate these analyses can be found at https://github.com/slrosen/reproductive_acceleration_lht.

## 9 GenAI use statement

We used Claude Code to help translate Stata code to R, clean and debug code, insert equation, figure, and section cross-referencing, create an outline for the supplementary materials, clean and alphabetize our .bib file, copy edit the text, create our README file, and assist with manuscript version control in GitHub.

1 Some empirical studies also separately test a looser version of this prediction by regressing *LRS* directly on phenotype with no interaction term, and thus asking whether acceleration has fitness benefits on average. This is a weaker claim than the hypothesis requires. The adaptive argument is that acceleration is particularly beneficial for individuals who experienced adversity, not that it is universally beneficial.

2 Weibel and colleagues reported qualitatively unchanged results when *LRS* was alternatively defined as the number of offspring that survived to 70 weeks, the average age at weaning.

3 Due to the field’s considerable interest in the effects of cumulative adversity, where adversity is operationalized as the total number of sources of adversity an organism experienced when they were young, we also provide parallel analyses using each baboon’s cumulative adversity count (ranging from 0–3+) as an alternative measure of early life adversity. These can be found in Section S3 of the supplementary materials. Results are qualitatively very similar to those reported for rainfall in the main text.

## References

Aduma, Mildred M, Gilbert O Ouma, Mohamed Y Said, Gordon O Wayumba, and Joseph Muhwang. 2018. “Spatial and Temporal Trends of Rainfall and Temperature in the Amboseli Ecosystem of Kenya.” Technical University of Kenya Institutional Repository 5 (5): 28–42.

Alberts, Susan C., and Jeanne Altmann. 2012. “The Amboseli Baboon Research Project: 40 Years of Continuity and Change.” In Long-Term Field Studies of Primates, edited by Peter M. Kappeler and David P. Watts. Springer Berlin Heidelberg. 10.1007/978-3-642-22514-7_12.

Altmann, Stuart A. 1998. Foraging for Survival: Yearling Baboons in Africa. University of Chicago Press.

André, Jean-Baptiste, and François Rousset. 2020. “Does Extrinsic Mortality Accelerate the Pace of Life? A Bare-Bones Approach.” Evolution and Human Behavior 41 (6): 486–92. 10.1016/j.evolhumbehav.2020.03.002.

Auld, Josh R., Anurag A. Agrawal, and Rick A. Relyea. 2009. “Re-Evaluating the Costs and Limits of Adaptive Phenotypic Plasticity.” Proceedings of the Royal Society B: Biological Sciences 277 (1681): 503–11. 10.1098/rspb.2009.1355.

Bateson, Patrick, Peter Gluckman, and Mark Hanson. 2014. “The Biology of Developmental Plasticity and the Predictive Adaptive Response Hypothesis.” The Journal of Physiology 592 (11): 2357–68. 10.1113/jphysiol.2014.271460.

Beldade, Patrícia, Ana Rita A. Mateus, and Roberto A. Keller. 2011. “Evolution and Molecular Mechanisms of Adaptive Developmental Plasticity.” Molecular Ecology 20 (7): 1347–63. 10.1111/j.1365-294X.2011.05016.x.

Belsky, Jay. 2012. “The Development of Human Reproductive Strategies: Progress and Prospects.” Current Directions in Psychological Science 21 (5): 310–16. 10.1177/0963721412453588.

Belsky, Jay, Paula L Ruttle, W Thomas Boyce, Jeffrey M Armstrong, and Marilyn J Essex. 2015. “Early Adversity, Elevated Stress Physiology, Accelerated Sexual Maturation, and Poor Health in Females.” Developmental Psychology 51 (6): 816.

Belsky, Jay, Laurence Steinberg, and Patricia Draper. 1991. “Childhood Experience, Interpersonal Development, and Reproductive Strategy: An Evolutionary Theory of Socialization.” Child Development 62 (4): 647–70. 10.1111/j.1467-8624.1991.tb01558.x.

Bonapersona, V, J Kentrop, CJ Van Lissa, R Van Der Veen, M Joëls, and RA Sarabdjitsingh. 2019. “The Behavioral Phenotype of Early Life Adversity: A 3-Level Meta-Analysis of Rodent Studies.” Neuroscience & Biobehavioral Reviews 102: 299–307.

Chisholm, James S, Peter T Ellison, Jeremy Evans, et al. 1993. “Death, Hope, and Sex: Life-History Theory and the Development of Reproductive Strategies [and Comments and Reply].” Current Anthropology 34 (1): 1–24.

Dantzer, Ben, Stan Boutin, Jeffrey E. Lane, and Andrew G. McAdam. 2022. “Integrative Studies of the Effects of Mothers on Offspring: An Example from Wild North American Red Squirrels.” In Patterns of Parental Behavior: From Animal Science to Comparative Ethology and Neuroscience, edited by Gabriela González-Mariscal. Springer International Publishing. 10.1007/978-3-030-97762-7_9.

Del Giudice, Marco. 2020. “Rethinking the Fast-Slow Continuum of Individual Differences.” Evolution and Human Behavior 41 (6): 536–49. 10.1016/j.evolhumbehav.2020.05.004.

Ding, Wenqin, Yuxiang Xu, Anthony J. Kondracki, and Ying Sun. 2024. “Childhood Adversity and Accelerated Reproductive Events: A Systematic Review and Meta-Analysis.” American Journal of Obstetrics and Gynecology 230 (3): 315–329.e31. 10.1016/j.ajog.2023.10.005.

Draper, Patricia, and Henry Harpending. 1982. “Father Absence and Reproductive Strategy: An Evolutionary Perspective.” Journal of Anthropological Research 38 (3): 255–73.

Ellis, Bruce J. 2004. “Timing of Pubertal Maturation in Girls: An Integrated Life History Approach.” Psychological Bulletin 130 (6): 920.

Ellis, Bruce J., Aurelio José Figueredo Barbara H. Brumbach, and Gabriel L. Schlomer. 2009. “Fundamental Dimensions of Environmental Risk.” Human Nature 20 (2): 204–68. 10.1007/s12110-009-9063-7.

Ellis, Samuel, Daniel W Franks, Stuart Nattrass, et al. 2018. “Postreproductive Lifespans Are Rare in Mammals.” Ecology and Evolution 8 (5): 2482–94. 10.1002/ece3.3856.

Frankenhuis, Willem E., and Alison Gopnik. 2023. “Early Adversity and the Development of Explore-Exploit Tradeoffs.” Trends in Cognitive Sciences 27 (7): 616–30. 10.1016/j.tics.2023.04.001.

Ghalambor, C. K., J. K. McKay, S. P. Carroll, and D. N. Reznick. 2007. “Adaptive Versus Non-Adaptive Phenotypic Plasticity and the Potential for Contemporary Adaptation in New Environments.” Functional Ecology 21 (3): 394–407. 10.1111/j.1365-2435.2007.01283.x.

Hayward, Adam D., Jari Holopainen, Jenni E. Pettay, and Virpi Lummaa. 2012. “Food and Fitness: Associations Between Crop Yields and Life-History Traits in a Longitudinally Monitored Pre-Industrial Human Population.” Proceedings of the Royal Society B: Biological Sciences 279 (1745): 4165–73. 10.1098/rspb.2012.1190.

Kuzawa, Christopher, Stacy Rosenbaum, and Lee Gettler. 2025. “No Need for Tiers to Explain the Ecological Predictors of Human Life History Strategies.” Behavioral and Brain Sciences 48: e112. 10.1017/S0140525X25100848.

Lafuente, Elvira, and Patrícia Beldade. 2019. “Genomics of Developmental Plasticity in Animals.” Frontiers in Genetics 10: 720. 10.3389/fgene.2019.00720.

Malani, Anup, Elizabeth A. Archie, and Stacy Rosenbaum. 2023. “Conceptual and Analytical Approaches for Modelling the Developmental Origins of Inequality.” Philosophical Transactions of the Royal Society B: Biological Sciences 378 (1883). 10.1098/rstb.2022.0306.

Monaghan, Pat. 2008. “Early Growth Conditions, Phenotypic Development and Environmental Change.” Philosophical Transactions of the Royal Society B: Biological Sciences 363 (1497): 1635–45. 10.1098/rstb.2007.0011.

Moorad, Jacob, Daniel Promislow, and Jonathan Silvertown. 2019. “Evolutionary Ecology of Senescence and a Reassessment of Williams’ ‘Extrinsic Mortality’ Hypothesis.” Trends in Ecology & Evolution 34 (6): 519–30. 10.1016/j.tree.2019.02.006.

Nettle, Daniel, and Melissa Bateson. 2015. “Adaptive Developmental Plasticity: What Is It, How Can We Recognize It and When Can It Evolve?” Proceedings of the Royal Society B: Biological Sciences 282 (1812): 20151005. 10.1098/rspb.2015.1005.

Parker, G. A., and J. Maynard Smith. 1990. “Optimality Theory in Evolutionary Biology.” Nature 348 (6296): 27–33. 10.1038/348027a0.

Petrullo, Lauren, David Delaney, Stan Boutin, Jeffrey E. Lane, Andrew G. McAdam, and Ben Dantzer. 2024. “A Future Food Boom Rescues the Negative Effects of Early-Life Adversity on Adult Lifespan in a Small Mammal.” Proceedings of the Royal Society B: Biological Sciences 291 (2021). 10.1098/rspb.2023.2681.

Pigliucci, Massimo. 2001. Phenotypic Plasticity: Beyond Nature and Nurture. Johns Hopkins University Press.

Rosenbaum, Stacy, Anup Malani, Amanda J. Lea, Jenny Tung, Susan C. Alberts, and Elizabeth A. Archie. 2025. “Testing Early Life Effects Frameworks: Developmental Constraints and Adaptive Response Hypotheses Do Not Explain Fertility Outcomes in Wild Female Baboons.” Proceedings of the Royal Society B: Biological Sciences 292 (2050). 10.1098/rspb.2024.2485.

Siracusa, Erin R, Xavier Bal, Delphine De Moor, et al. 2025. “Environment Dependent Benefits of Sociality in Soay Sheep.” bioRxiv, 2025.05.04.652099. 10.1101/2025.05.04.652099.

Snyder-Mackler, Noah, Joseph Robert Burger, Lauren Gaydosh, et al. 2020. “Social Determinants of Health and Survival in Humans and Other Animals.” Science 368 (6493): eaax9553. 10.1126/science.aax9553.

Stearns, Stephen C. 1992. The Evolution of Life Histories. Oxford University Press. 10.1093/oso/9780198577416.001.0001.

Tung, Jenny, Elizabeth A. Archie, Jeanne Altmann, and Susan C. Alberts. 2016. “Cumulative Early Life Adversity Predicts Longevity in Wild Baboons.” Nature Communications 7 (1): 11181. 10.1038/ncomms11181.

Uller, Tobias. 2008. “Developmental Plasticity and the Evolution of Parental Effects.” Trends in Ecology & Evolution 23 (8): 432–38. 10.1016/j.tree.2008.04.005.

Warner, Daniel A. 2014. “Fitness Consequences of Maternal and Embryonic Responses to Environmental Variation: Using Reptiles as Models for Studies of Developmental Plasticity.” Integrative and Comparative Biology 54 (5): 757–73. 10.1093/icb/icu099.

Weibel, Chelsea J., Jenny Tung, Susan C. Alberts, and Elizabeth A. Archie. 2020. “Accelerated Reproduction Is Not an Adaptive Response to Early-Life Adversity in Wild Baboons.” Proceedings of the National Academy of Sciences 117 (40): 24909–19. 10.1073/pnas.2004018117.

West-Eberhard, Mary Jane. 2003. Developmental Plasticity and Evolution. Oxford University Press.

Zipple, Matthew N., Jeanne Altmann, Fernando A. Campos, et al. 2021. “Maternal Death and Offspring Fitness in Multiple Wild Primates.” Proceedings of the National Academy of Sciences 118 (1): e2015317118. 10.1073/pnas.2015317118.

Zipple, Matthew N., H. Kern Reeve, and Orca Jimmy Peniston. 2024. “Maternal Care Leads to the Evolution of Long, Slow Lives.” Proceedings of the National Academy of Sciences 121 (25): e2403491121. 10.1073/pnas.2403491121.

Zipple, Matthew N., Caleb C. Vogt, and Michael J. Sheehan. 2024. “Genetically Identical Mice Express Alternative Reproductive Tactics Depending on Social Conditions in the Field.” Proceedings of the Royal Society B: Biological Sciences 291 (2019). 10.1098/rspb.2024.0099.

