## Supplementary materials for "Accounting for the life history structure of fitness in tests for adaptive reproductive acceleration"

Accepted at *Philosophical Transactions of the Royal Society B.*
10.1098/rstb.2025.0479

Stacy Rosenbaum & Anup Malani

**4 Omission of control variables in the reproductive pacing and adversity regressions** **9**

**5 Full Taylor expansion regression results** **10**

**References** **11**

### 1 Amboseli baboon data details

The Amboseli Baboon Research Project (ABRP) has monitored wild baboons living in the Amboseli ecosystem of southern Kenya since 1971 (Alberts and Altmann 2012). The population is made up of yellow baboons (*Papio cynocephalus*) with anubis (*P. anubis*) admixture from a neighboring, also wild population (Wall et al. 2016). Each animal is individually identifiable by trained observers, who collect demographic and behavioral data on each social group multiple times per week year-round. This allows the project to capture fine-grained data on life history events, including births, infant loss, lifespans, and other metrics necessary for the analyses presented in this paper (Alberts and Altmann 2012). Here we only use female data because it is much harder to measure reproductive pacing in males, and because male baboons disperse at sexual maturity and thus longitudinal data across adulthood are not typically available.

Most of the data used here were previously published by Weibel and colleagues (2020). Briefly, lifetime reproductive success was defined as the total number of live offspring the subject female gave birth to, and age at first birth was defined as her age (in years) when she gave birth to her first live offspring. Interbirth intervals were defined as the number of days between consecutive live births; to count as two consecutive live births, the first offspring in the series needed to have survived to at least 70 weeks, which is approximately the age at which baboons are weaned (Altmann 1998). The original paper (and thus this paper) excluded females whose measure (either age at first birth or interbirth interval) was greater than three standard deviations from the mean, in order to eliminate females with potential pathological problems. These exclusions represented a tiny fraction of the total number of measures (1.7% of possible age at first birth measurements, and 0.01% of individual IBIs). More details about the original data analysis, including data collection and summary methodology, can be found in Weibel et al. (2020).

The one addition we made to the original data was moving from a categorical version of the rainfall variable (drought/no drought; see the section below for more details) to a continuous version. This is better-suited to our analyses for two reasons. First, it preserves information about the full range of rainfall variation the baboons actually experience. Second, the Taylor expansion regression requires a continuous predictor to generate meaningful squared and interaction terms. With categorical predictors, these terms collapse; e.g., a 0/1 variable squared is still 0/1, so the coefficients are not interpretable.

The ABRP team supplied us with the continuous rainfall data underlying the categorical data used in the original paper. These data are collected via daily monitoring of a rain gauge at the ABRP camp. Across the 267 females who appear in the data set, mean cumulative rainfall during the first year of life was 321mm (SD = 122mm), and ranged from 92mm to 767mm. These are very similar to the values reported for a slightly larger sample of females (n=295) in another recent Amboseli paper that relied on first-year-of-life rainfall data (Rosenbaum et al. 2025), along with longitudinal precipitation data for Amboseli across time (Southworth et al. 2026).

#### 2 Scope of the accounting and optimization models

##### 2.1 Extension of the accounting model to females with lifetime reproductive success <2

The accounting model in Equation 4 of the main text contains an average interbirth interval ($IBI$) term, which is undefined for females with only a single birth. By construction, the model applies only to females with $LRS \geq 2$. Thus, females with $LRS = 0$ or $LRS = 1$ are not included in Figure 3 in the main text.

However, the model could be extended to females with $LRS = 1$ by substituting a population-mean $IBI$ for the missing observed value. Starting from $L \approx A + (LRS - 0.5) \cdot IBI$, the rearranged form $LRS \approx (L - A)/IBI_{\text{pop}} + 0.5$ gives us an unbiased prediction *on average*, because the model's key assumption is that the time of survival after the final birth ($E$) averages $0.5 \cdot IBI$. Individual predictions will have a larger relative error than for $LRS \geq 2$ females, for two reasons. First, assigning the population-mean $IBI$ rather than the female's own (unknown) individual $IBI$ adds imputation noise that $LRS \geq 2$ females are not subject to. Second, the same absolute prediction error from individual deviation around the model's central assumption ($E \approx 0.5 \cdot IBI$) has the largest relative impact at $LRS = 1$, because $L - A$ is at its smallest there ($0.5 \cdot IBI$). This means that any given error is a larger fraction of $L - A$ itself. A female who dies shortly after her one birth receives a predicted $LRS$ of about 0.5, while a female who survives nearly a full $IBI$ without a second birth receives a prediction closer to 1.5. This is a much wider relative error than the analogous range at higher $LRS$: e.g., at $LRS = 5$, the deviation changes the prediction from 4.5 to 5.5, which is only 20% of the target $LRS = 5$, instead of 100% when $LRS = 1$. The extension is therefore feasible, but would be expected to introduce substantial variance at the individual level.

To test how the accounting model performs with an imputed $IBI$ for females where $LRS = 1$, we returned to the Weibel et al. (2020) data, which contains 19 females with $LRS = 1$. Applying the rearranged equation with the population-mean $IBI$ (678 days), the mean predicted $LRS$ for these females is 1.02 (expected: 1.00), confirming that the extension to this additional subset of females is unbiased on average. Individual predictions range from 0.50 to 1.88, with a standard deviation of 0.39 (Figure S1).

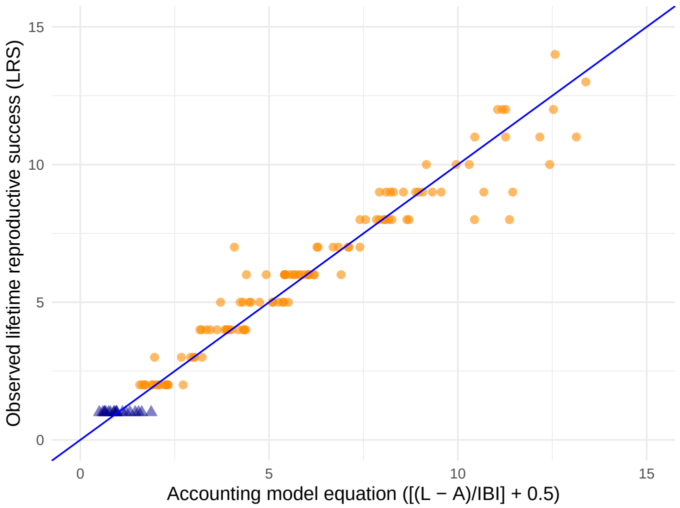

Figure S1: Empirical illustration of the $LRS = 1$ extension of the accounting model. Orange circles show females with $LRS \geq 2$ (n = 110, which is the same data subset used in Figure 3 of the main text). Blue triangles show values for the 19 females with $LRS = 1$, for whom predicted $LRS$ was computed using the rearranged accounting equation with a population-mean $IBI$. The blue line is the 1:1 fit line. The blue triangles cluster around an observed $LRS$ of 1 (by definition) with predicted values centered near 1 and spanning both sides of the fit line. This illustrates that the extension is unbiased on average but noisy for individual females.

We also repeated the regression-based validation from §2.1 of the main text on this larger dataset. Table S1 reports the same two specifications (linear and log forms of the accounting model) used in Table 1 of the main text, but using the combined sample of females with $LRS \geq 2$ and $LRS = 1$. In the linear version, the explained variance is the same as in Table 1 ($R^2 = 0.98$ in both samples), and the estimated coefficients on $L$ and $A$ are slightly closer to the values the accounting equation predicts (1 and $-1$, respectively). Unsurprisingly, the log specification does not perform as well: $R^2$ drops from 0.99 to 0.92. This is because for females where $LRS = 1$ both $\log(LRS - 0.5)$ and the imputed $\log(IBI)$ are constants that have only a single value. Thus, most of the within-group variation that the log specification relies on is absent. This is an expected cost of data imputation, rather than a problem with the accounting model itself.

**Table S1:** Accounting regression results for extended sample including females where LRS = 1.

| | (LRS-0.5)*IBI | log(LRS-0.5) |
|---|---|---|
| L (Lifespan) | 0.938*** (0.022) | |
| A (Age at first birth) | −0.825*** (0.058) | |
| log(L - A) | | 0.755*** (0.030) |
| log(IBI) | | −0.182* (0.102) |
| R2 | 0.981 | 0.923 |
| Num.Obs. | 129 | 129 |

Standard errors in parentheses.

* p < 0.10, ** p < 0.05, *** p < 0.01

In contrast to females with $LRS = 1$, females with $LRS = 0$ are outside the scope of the reproductive acceleration hypothesis itself, not just outside the model's mechanical domain (though they are also outside of that, given that they have neither an age at first birth nor an $IBI$). The hypothesis asks whether females who experience early adversity can improve their fitness by accelerating their reproductive schedule, which is only a relevant question for females who actually reproduce. Females who die before reproducing have zero direct fitness regardless of what reproductive schedule they would (hypothetically) have adopted. Thus, their absence from the model does not represent missing evidence about reproductive acceleration. If early adversity increases pre-reproductive mortality, leading to $LRS = 0$, that pattern would provide evidence for a different class of hypotheses: developmental constraints models, including the "silver spoon" hypothesis, which propose that early adversity compromises organisms in ways that cannot be compensated for later (Lea and Rosenbaum 2020; Malani et al. 2023).

#### 2.2 Generalization of the accounting model across species

The accounting model generalizes to many other iteroparous species. Its assumption is that post-final-birth survival is half of a typical $IBI$, and there is no compelling reason to believe that this uniquely applies to baboons. In species where females produce multiple offspring per reproductive event, the most natural extension would be to redefine $LRS$ as the number of reproductive events (i.e., litters). While it could theoretically still use total offspring count instead, it would require that litter size be roughly stable across a female's lifetime. Important exceptions are humans and the small number of other mammalian species with substantial post-reproductive lifespans, such as killer whales and short-finned pilot whales (Johnstone and Cant 2019). For these species the assumption that females tend to die roughly mid-reproductive cycle breaks down, because they stop reproducing well before they die. However, in theory the model could be extended to such species if we had dependable estimates of the average post-reproductive lifespan to substitute for the $0.5 \cdot IBI$ approximation.

#### 2.3 Generalization of the optimization model within species

A separate scope condition applies to the optimization model rather than to the accounting model. The optimization model's predictions about how the optimum shifts with adversity (Equation 8) assume that there is a single common trade-off function $L(A, IBI, e)$ across females, where $e$ is some feature of the early life environment. In a hypothetical world where different individuals within populations have stable alternative reproductive tactics — i.e., distinct phenotypes with different underlying trade-off functions — these predictions would apply within each phenotype rather than across the population as a whole. We do not have evidence for such tactics in the Amboseli baboons, but this would be an important modification to consider when applying these methods to species where stable alternative tactics exist.

### 3 Cumulative adversity in the Amboseli baboons

While we focus specifically on rainfall during the first year of life in the main text, this is certainly not the only source of hardship young baboons experience. The Amboseli Baboon Research Project has identified six sources of early adversity for its study animals, which are described in detail in Table S2. Many prior papers on the baboons have examined the effects of these sources cumulatively (e.g., Tung et al. 2016; Weibel et al. 2020; Rosenbaum et al. 2020; Anderson et al. 2024), where each source is coded as a yes/no indicator (the animal in question did/did not experience that source). A baboon's cumulative adversity count is the number of "yes" responses, but because few baboons experience more than three sources, counts of 3+ are collapsed into a single category. This cumulative adversity approach is standard across the broader literature (reviewed for human studies in McLaughlin and Sheridan 2016).

**Table S2:** Sources of early life adversity for Amboseli baboons.

| Source of adversity | Definition (from Tung et al. 2016) |
|---|---|
| Drought | <200mm of cumulative rainfall during the subject's first year of life. |
| High population density | Number of adults of both sexes in the subject's group on the day the subject was born; coded as "yes" if group size fell in the largest quartile of the population distribution. |
| Low maternal dominance rank | Ordinal dominance rank of the subject's mother in the month that the subject was born; coded as "yes" if the mother's rank fell in the lowest quartile of the population distribution. |
| Low maternal social connectedness | Social connectedness (measured by grooming relationships) of a subject's mother to other adult females, calculated as the subject's mother's average connectedness value over the first two years of the subject's life. Coded as "yes" if the subject's mother fell in the lowest quartile of the population. |
| Early maternal loss | Death of the subject's mother before the subject turns 4 years old. |
| Competing younger sibling | The subject had a sibling born less than 1.5 years after the subject was born; 1.5 years corresponds to the lowest quartile of interbirth intervals in the baboon population. |

#### 3.1 The reproduction–lifespan trade-off

Here we present the regression results for the reproduction–lifespan trade-off analyses (i.e., the Taylor expansion regressions) but replace rainfall (our environmental variable, $e$) with cumulative early life adversity (Table S3; see Section 5.1.1 in the main text). Table S3 contains results with both lifespan and the rearranged accounting model $(LRS - 0.5) \cdot IBI$ as the outcome variable. (The latter is equivalent to asking about the pacing–fitness trade-off.) Note that cumulative adversity is coded in the opposite direction from rainfall. Higher values indicate *more* adversity, so the predicted signs on $e$ and its interactions flip: under the reproductive acceleration hypothesis, coefficients from the regressions of $A$ on $e$ and $IBI$ on $e$, as well as $\lambda_{A \cdot e}$ and $\lambda_{IBI \cdot e}$, should all be negative rather than positive. Sample sizes are smaller in these analyses (n=61, versus n=99 in the main text) because not all females have complete cumulative adversity data.

Our conclusions are robust to this alternative choice of adversity measure. We again find no evidence for the reproductive acceleration hypothesis. The regression explains a moderate amount of lifespan variation ($R^2 = 0.25$), but the key interaction coefficients $\lambda_{A \cdot e}$ and $\lambda_{IBI \cdot e}$ are not statistically significant ($p > 0.30$ for both; see Table S3, first column). Furthermore, the point estimates run in the opposite direction of the hypothesis prediction, suggesting that if anything, adversity steepens rather than flattens the reproduction–lifespan trade-off. We find a similar null result when we use $(LRS - 0.5) \cdot IBI$ as the outcome (second column of Table S3). The interaction coefficients are again non-significant ($p > 0.27$).

**Table S3:** Taylor expansion regression results using cumulative adversity

| | Lifespan | (LRS − 0.5) · IBI |
|---|---|---|
| (Intercept) | 42.494 (56.494) | 49.209 (54.189) |
| A | −10.878 (22.952) | −13.636 (22.015) |
| IBI | 19.817 (29.203) | 14.325 (28.011) |
| Cumul. adversity | −11.491 (9.767) | −8.392 (9.369) |
| $A^2$ | 0.009 (2.122) | 0.004 (2.035) |
| $IBI^2$ | −13.484** (5.577) | −12.615** (5.349) |
| Cumul. adversity$^2$ | −0.429 (0.885) | −0.591 (0.849) |
| A × IBI | 4.619 (3.827) | 5.604 (3.670) |
| A × Cumul. adv. | 1.689 (1.628) | 1.712 (1.562) |
| IBI × Cumul. adv. | 0.340 (2.957) | −1.176 (2.836) |
| R2 | 0.252 | 0.244 |
| R2 Adj. | 0.119 | 0.111 |
| Num.Obs. | 61 | 61 |

Standard errors in parentheses.

* p < 0.10, ** p < 0.05, *** p < 0.01

In the second column, the coefficient on A is the Taylor expansion coefficient minus 1 (because the accounting substitution subtracts A from both sides).

##### 3.2 Reproductive pacing and adversity

Here we execute the reproductive pacing analyses from Section 5.1.2 of the main text, but we replace rainfall with cumulative adversity (Table S4). As described above, cumulative adversity is coded in the opposite direction from rainfall (higher values indicate more adversity), so if the reproductive acceleration hypothesis is supported we would expect negative coefficients: more adversity should lead to earlier first births and shorter IBIs.

We again find no evidence for reproductive acceleration. Cumulative adversity does not significantly predict age at first birth ($p = 0.19$) or interbirth interval ($p = 0.44$), and the point estimates for both outcomes run in the opposite direction from the hypothesis prediction.

**Table S4:** Reproductive pacing regression results for cumulative adversity

| | Age at first birth | Interbirth interval |
|---|---|---|
| (Intercept) | 6.143*** (0.076) | 1.839*** (0.076) |
| Cumul. adversity | 0.062 (0.047) | 0.041 (0.053) |
| R2 | 0.008 | 0.010 |
| Num.Obs. | 211 | 61 |

Standard errors in parentheses.

* p < 0.10, ** p < 0.05, *** p < 0.01

### 4 Omission of control variables in the reproductive pacing and adversity regressions

Weibel and colleagues' (2020) analysis included two control variables in models of reproductive pacing: the population growth rate during the focal female's infancy (which captures population-level resource availability during early life) and the number of mature females in the group at the time of first birth (a proxy for within-group competition for resources at the onset of reproduction). We opted not to include these controls in our models for two reasons. First, they are not motivated by the optimization model we derive in the main text; including them would add complexity without a clear theoretical rationale. Second, population growth rate is conceptually related to our rainfall adversity measure, since both reflect resource availability, which could make interpretation less straightforward (though in practice the two are only weakly correlated: $r = 0.07$). To verify that this choice does not affect our conclusions, we re-ran all of the reproductive pacing and adversity regressions, from both the main text and the reproductive pacing and adversity section above, with both controls included. The adversity coefficients (either rainfall or cumulative adversity) are very similar in magnitude, sign, and significance whether or not controls are included. We therefore report the simpler models without controls in the main text.

**Table S5:** Reproductive pacing regression results with and without control variables for age at first birth

| | Rainfall | Rainfall + ctrls | Cumul. adv. | Cumul. adv. + ctrls |
|---|---|---|---|---|
| (Intercept) | 6.510*** (0.110) | 6.696*** (0.205) | 6.143*** (0.076) | 6.475*** (0.250) |
| Rainfall (first year) | −0.001*** (0.000) | −0.001*** (0.000) | | |
| Pop growth (×1000) | | −0.276* (0.144) | | −0.318 (0.195) |
| Group size (females) | | 0.005 (0.006) | | 0.000 (0.007) |
| Cumul. adversity | | | 0.062 (0.047) | 0.058 (0.047) |
| R2 | 0.037 | 0.052 | 0.008 | 0.021 |
| Num.Obs. | 267 | 267 | 211 | 211 |

**Table S6:** Reproductive pacing regression results with and without control variables for interbirth intervals

| | Rainfall | Rainfall + ctrls | Cumul. adv. | Cumul. adv. + ctrls |
|---|---|---|---|---|
| (Intercept) | 1.944*** (0.090) | 1.831*** (0.172) | 1.839*** (0.076) | 1.875*** (0.209) |
| Rainfall (first year) | 0.000 (0.000) | 0.000 (0.000) | | |
| Pop growth (×1000) | | −0.054 (0.089) | | −0.142 (0.152) |
| Group size (females) | | 0.009 (0.006) | | 0.006 (0.007) |
| Cumul. adversity | | | 0.041 (0.053) | 0.053 (0.054) |
| R2 | 0.017 | 0.047 | 0.010 | 0.036 |
| Num.Obs. | 99 | 99 | 61 | 61 |

Standard errors in parentheses.

* $p < 0.10$, ** $p < 0.05$, *** $p < 0.01$

Population growth rate rescaled (×1000) from original units.

### 5 Full Taylor expansion regression results

Table S7 shows full Taylor expansion regression results when rainfall is used as the measure of early life adversity. This corresponds to the summary results reported in Section 5.1.1 of the main text. In the first column, lifespan is the outcome; in the second column, the outcome is the rearranged version of the accounting model $(LRS - 0.5) \cdot IBI$, which answers questions about reproductive pacing–fitness trade-offs. The third column reports a simple regression $L \sim e$ with no reproductive pacing variables included, which captures the total relationship between rainfall and lifespan. We report it for transparency and as the diagnostic discussed in the main text. The rainfall coefficient has the same sign as in the full Taylor expansion, indicating that including $A$ and $IBI$ in the structural model is not introducing the kind of bias that would arise if unobserved factors were jointly influencing both pacing and lifespan.

**Table S7:** Full Taylor expansion regression results using rainfall as the measure of early life adversity.

| | Lifespan (full) | (LRS − 0.5) · IBI | Lifespan (simple) |
|---|---|---|---|
| (Intercept) | −25.689 (48.954) | −13.735 (47.760) | 14.303*** (1.432) |
| A | 14.446 (15.281) | 11.543 (14.908) | |
| IBI | −6.145 (20.618) | −13.966 (20.115) | |
| Rainfall | 0.022 (0.069) | 0.016 (0.067) | 0.006 (0.004) |
| $A^2$ | −1.840 (1.327) | −1.844 (1.294) | |
| $IBI^2$ | −9.964** (4.814) | −8.998* (4.697) | |
| $Rainfall^2$ | 0.000 (0.000) | 0.000 (0.000) | |
| A × IBI | 6.058* (3.079) | 7.022** (3.004) | |
| A × Rainfall | −0.008 (0.012) | −0.007 (0.012) | |
| IBI × Rainfall | 0.015 (0.019) | 0.014 (0.019) | |
| R2 | 0.117 | 0.133 | 0.021 |
| R2 Adj. | 0.028 | 0.045 | 0.011 |
| Num.Obs. | 99 | 99 | 99 |

Standard errors in parentheses.

* $p < 0.10$, ** $p < 0.05$, *** $p < 0.01$

In the second column, the coefficient on A is the Taylor expansion coefficient minus 1 (because the accounting substitution subtracts A from both sides).

### References

Alberts, Susan C., and Jeanne Altmann. 2012. “The Amboseli Baboon Research Project: 40 Years of Continuity and Change.” In *Long-Term Field Studies of Primates*, edited by Peter M. Kappeler and David P. Watts. Springer Berlin Heidelberg. https://doi.org/10.1007/978-3-642-22514-7_12.

Altmann, Stuart A. 1998. *Foraging for Survival: Yearling Baboons in Africa*. University of Chicago Press.

Anderson, Jordan A., Dana Lin, Amanda J. Lea, et al. 2024. “DNA Methylation Signatures of Early-Life Adversity Are Exposure-Dependent in Wild Baboons.” *Proceedings of the National Academy of Sciences* 121 (11): e2309469121. https://doi.org/10.1073/pnas.2309469121.

Johnstone, Rufus A., and Michael A. Cant. 2019. “Evolution of Menopause.” *Current Biology* 29 (4): R112–15. https://doi.org/10.1016/j.cub.2018.12.048.

Lea, Amanda J., and Stacy Rosenbaum. 2020. “Understanding How Early Life Effects Evolve: Progress, Gaps, and Future Directions.” *Current Opinion in Behavioral Sciences* 36: 29–35. https://doi.org/10.1016/j.cobeha.2020.06.006.

Malani, Anup, Elizabeth A. Archie, and Stacy Rosenbaum. 2023. “Conceptual and Analytical Approaches for Modelling the Developmental Origins of Inequality.” *Philosophical Transactions of the Royal Society B: Biological Sciences* 378 (1883). https://doi.org/10.1098/rstb.2022.0306.

McLaughlin, Katie A, and Margaret A Sheridan. 2016. “Beyond Cumulative Risk: A Dimensional Approach to Childhood Adversity.” *Current Directions in Psychological Science* 25 (4): 239–45. https://doi.org/10.1177/096372141665.

Rosenbaum, Stacy, Anup Malani, Amanda J. Lea, Jenny Tung, Susan C. Alberts, and Elizabeth A. Archie. 2025. “Testing Early Life Effects Frameworks: Developmental Constraints and Adaptive Response Hypotheses Do Not Explain Fertility Outcomes in Wild Female Baboons.” *Proceedings of the Royal Society B: Biological Sciences* 292 (2050). https://doi.org/10.1098/rspb.2024.2485.

Rosenbaum, Stacy, Shuxi Zeng, Fernando A. Campos, et al. 2020. “Social Bonds Do Not Mediate the Relationship Between Early Adversity and Adult Glucocorticoids in Wild Baboons.” *Proceedings of the National Academy of Sciences* 117 (33): 20052–62. https://doi.org/10.1073/pnas.2004524117.

Southworth, Chelsea A., Jack C. Winans, Jacob B. Gordon, et al. 2026. “Demographic, Behavioral, and Ecological Data from a Long-Term Field Study of Wild Baboons in Amboseli, Kenya.” *Scientific Data* 13 (1): 311. https://doi.org/10.1038/s41597-026-06741-2.

Tung, Jenny, Elizabeth A. Archie, Jeanne Altmann, and Susan C. Alberts. 2016. “Cumulative Early Life Adversity Predicts Longevity in Wild Baboons.” *Nature Communications* 7 (1): 11181. https://doi.org/10.1038/ncomms11181.

Wall, Jeffrey D, Stephen A Schlebusch, Susan C Alberts, et al. 2016. “Genomewide Ancestry and Divergence Patterns from Low-coverage Sequencing Data Reveal a Complex History of Admixture in Wild Baboons.” *Molecular Ecology* 25 (14): 3469–83. https://doi.org/10.1111/mec.13684.

Weibel, Chelsea J., Jenny Tung, Susan C. Alberts, and Elizabeth A. Archie. 2020. “Accelerated Reproduction Is Not an Adaptive Response to Early-Life Adversity in Wild Baboons.” *Proceedings of the National Academy of Sciences* 117 (40): 24909–19. https://doi.org/10.1073/pnas.2004018117.
